# Astroglial FMRP modulates synaptic signaling and behavior phenotypes in FXS mouse model

**DOI:** 10.1101/2020.02.10.941971

**Authors:** Shan-Xue Jin, Haruki Higashimori, Christina Schin, Alessandra Tamashiro, Yuqin Men, Ming Sum R Chiang, Rachel Jarvis, Dan Cox, Larry Feig, Yongjie Yang

## Abstract

Fragile X syndrome (FXS) is one of the most common inherited intellectual disability (ID) disorders, in which the loss of FMRP protein induces a range of cellular signaling changes primarily through excess protein synthesis. Although neuron-centered molecular and cellular events underlying FXS have been characterized, how different CNS cell types are involved in typical FXS synaptic signaling changes and behavioral phenotypes is largely unknown. Recent evidence suggests that selective loss of astroglial FMRP is able to dysregulate glutamate uptake, increase spine density, and impair motor-skill learning. Here we investigated the effect of astroglial FMRP on synaptic signaling and FXS-related behavioral and learning phenotypes in astroglial Fmr1 cKO and cON mice in which FMRP expression is selectively diminished or restored in astroglia. We found that selective loss of astroglial FMRP contributes to cortical hyperexcitability by enhancing NMDAR-mediated evoked but not spontaneous miniEPSCs and elongating cortical UP state duration. Selective loss of astroglial FMRP is also sufficient to increase locomotor hyperactivity, significantly diminish social novelty preference, and induce memory acquisition and extinction deficits in astroglial Fmr1 cKO mice. Importantly, re-expression of astroglial FMRP is able to significantly rescue the hyperactivity (evoked NMDAR response, UP state duration, and open field test) and social novelty preference in astroglial Fmr1 cON mice. These results demonstrate a profound role of astroglial FMRP in the evoked synaptic signaling, spontaneously occurring cortical UP states, and FXS-related behavioral and learning phenotypes and provide important new insights in the cell type consideration for the FMRP reactivation strategy.

## Introduction

Fragile X syndrome (FXS) is one of the most common inherited intellectual disabilities (ID), which manifests with cognitive impairment, hyperactivity/seizure, sensory hypersensitivity, and several autistic features such as repetitive behaviors and social withdraw (1). In FXS, the *fmr1* gene on the X chromosome is fully mutated with an expansion of more than 200 CGG repeats at its promoter and becomes hypermethylated (2), leading to the transcriptional silencing of the *fmr1* gene and loss of FMRP protein (3). FMRP has been demonstrated to be an RNA-binding protein that regulates translation of many (> 1000) mRNAs (3, 4). The loss of FMRP function generally leads to excess protein synthesis including the enhancement of several intracellular signaling pathways in neurons, including BDNF (5), PI3K (6, 7), GSK3β (8), and insulin (9), etc. By studying the mouse model (Fmr1 KO mice) of FXS (10), altered synaptic plasticity, particularly enhanced mGluR5-dependent long-term depression (LTD) in hippocampus (11, 12), has been identified as a major mechanism underlying the learning disability of FXS. Dysfunction of the GABA system, either reduced GABA receptor subunit expression (13) or an impaired developmental excitatory-to-inhibitory switch (14), has also been implicated in the hyperexcitability phenotype of FXS. In addition, a translation-independent function of FMRP, by directly binding with the BK channel’s β4 subunit, has also been identified that leads to elevated neurotransmitter release from pre-synaptic terminals and enhancement of neuronal excitability (15).

How the loss of FMRP in different brain regions/cell types specifically impacts FXS phenotypes has begun to shed new light on the region/cell specificity contributing to FXS pathophysiology. Selective deletion of the *fmr1* gene in Purkinje neurons leads to enhanced LTD and deficits in eye-blink conditioning (16). Complete deletion of FMRP in a large number (60%) of cortical and hippocampal neurons, however, only showed activated Akt-mTOR pathway signaling without apparent behavioral or synaptic phenotypes (17). Selective ablation of FMRP in adult neural stem cells leads to reduced hippocampal neurogenesis and disrupts hippocampus-dependent learning (18). Forebrain excitatory neuron-selective deletion of the *fmr1* gene is sufficient to enhance cortical MMP-9 gelatinase activity, mTOR/Akt phosphorylation, and resting EEG gamma power (19). By generating astroglial Fmr1 cKO and cON mice in which FMRP expression is selectively diminished or restored (20), we previously showed that selective loss of astroglial FMRP dysregulates astroglial glutamate transporter GLT1 expression which contributes to neuronal excitability as well as increased spine density (20). An independent study with a different set of astroglia specific Fmr1 KO mice also displayed an increased spine density in the motor cortex and impaired motor-skill learning (21). Despite these observations, how loss of astroglial FMRP affects synaptic signaling and behavioral phenotypes of FXS remains unexplored. To answer these questions, here we examined these phenotypes in astroglial Fmr1 cKO and cON mice. Our results demonstrate a profound role of astroglial FMRP in evoked synaptic signaling, spontaneously occurring cortical UP states, and FXS-related behavioral and learning phenotypes. These results also shed light on much understudied role of astroglia in neurodevelopmental disorders and intellectual disabilities.

## Results

### Selective diminishment of astroglial FMRP expression enhances NMDAR-mediated evoked responses but not spontaneous miniature excitatory post synaptic currents (mEPSCs)

Alterations of synaptic signaling mediated by different synaptic receptors in Fmr1 KO mice have been widely observed in multiple brain regions (cortex, hippocampus, cerebellum, etc.) (1, 22). Loss of FMRP leads to a dramatic switch of the NMDA/AMPA ratio during the postnatal P4 to P7 period in somatosensory cortex of Fmr1 KO mice (23), likely due to the retention of NMDAR-only immature silent synapses (23). However, increased AMPA/NMDA ratio, particularly reduced NMDAR currents were also found in either prefrontal or anterior piriform cortex of adult (aged) Fmr1 KO mice (24, 25). We previously showed that the selective loss of astroglial FMRP modestly increases spine density and length in somatosensory cortical pyramidal neurons of astroglial Fmr1 cKO mice, which can be rescued by the selective re-expression of FMRP in astroglia in astroglial Fmr1 cON mice (20). To investigate whether expression changes of FMRP solely in astroglia alters synaptic signaling of somatosensory cortical neurons, whole-cell voltage clamp recordings were performed with pyramidal neurons in layer 5 of somatosensory cortex at postnatal day (P) 38-45 from Fmr1 KO, astroglial Fmr1 cKO, and cON mice. The generation of astroglial Fmr1 cKO and cON mice using the astroglia-specific *slc1a3*-CreER driver line has been previously described (20) and the selective deletion or restoration of FMRP expression in astroglia in astroglial Fmr1 cKO (Cre^+^Fmr1^f/y^) and cON (Cre^+^Fmr1^loxP-neo/y^) mice has also been previously characterized (26). We have also bred astroglial Fmr1 cKO and cON mice with astroglial reporter Bac *aldh1l1*-eGFP mice so that cortical astroglia can be selectively isolated through fluorescence activated cell sorting (FACS)(27). Immunoblotting of FMRP from these acutely isolated cortical astroglia confirmed that FMRP levels are significantly reduced or restored in Cre^+^Fmr1^f/y^ or Cre^+^Fmr1^loxP-neo/y^ mice as expected (Fig. S1A). The properties of patched pyramidal neurons were confirmed with neuronal morphology (Fig. S1B) and firing patterns (Fig. S1C).

We first recorded miniature excitatory post synaptic currents (mEPSCs) by holding the membrane at −70 mV and pharmacologically isolating AMPAR or NMDAR-mediated currents respectively (more details in Materials and Methods). Unlike the well-observed reduced AMPAR activity in hippocampus (6, 28), no significant difference in the amplitude or frequency of AMPAR-mediated mEPSCs between neurons from Fmr1^+/y^ and Fmr1^-/y^ mice was observed (amplitude: Fmr1^+/y^, 24.73 ± 1.00 pA, n=17; Fmr1^-/y^, 28.22 ± 1.93 pA, n=13, p>0.05; frequency: Fmr1^+/y^, 5.58 ± 0.52 Hz, n=17; Fmr1^-/y^, 5.77 ± 0.65 Hz, n=13, p>0.05) (Fig. 1 A-C). We also found no differences in AMPAR mEPSC amplitude and frequency between pyramidal neurons from Cre^-^Fmr1^f/y^ (control) and Cre^+^Fmr1^f/y^ (cKO) mice in which the *fmr1* gene is selectively deleted in astroglia (amplitude: Cre^-^Fmr1^f/y^, 26.40 ± 1.13 pA, n=17; Cre^+^Fmr1^f/y^, 22.19 ± 1.15 pA, n=13, p>0.05; frequency: Cre^-^Fmr1^f/y^, 5.49 ± 0.49 Hz, n=17; Cre^+^Fmr1^f/y^, 5.15 ± 0.76 Hz, n=13, p>0.05) or from Cre^-^Fmr1^loxP-neo/y^ (control) and Cre^+^Fmr1^loxP-neo/y^ (cON) mice in which FMRP is selectively re-expressed in astroglia (amplitude: Cre^-^Fmr1^loxP-neo/y^, 26.84 ± 1.50 pA, n=15; Cre^+^Fmr1^loxP-neo/y^, 23.59 ± 0.65 pA, n=18, p>0.05; frequency: Cre^-^Fmr1^loxP-neo/y^, 5.47 ± 0.52 Hz, n=15; Cre^+^Fmr1^loxP-neo/y^, 6.25 ± 0.16 Hz, n=18, p>0.05) (Fig. 1 A-C). These results indicate that the loss of FMRP globally or selectively from astroglia has no detectable effect on AMPAR activity in pyramidal neurons of the somatosensory cortex. By contrast, our recordings found significant differences in NMDAR-mediated mEPSC amplitude but not frequency in pyramidal neurons between Fmr1^+/y^ and Fmr1^-/y^ mice (amplitude: Fmr1^+/y^, 4.51 ± 0.18 pA, n=20; Fmr1^-/y^, 6.05 ± 0.31 pA, n=24, p<0.0001; frequency: Fmr1^+/y^, 8.35 ± 0.41 Hz, n=20; Fmr1^-/y^, 8.04 ± 0.75 Hz, n=24, p>0.05)(Fig. 1 D-F). However, no differences in NMDAR-mediated mEPSC amplitude and frequency were found between neurons from Cre^-^Fmr1^f/y^ (amplitude: 5.21 ± 0.12 pA; frequency: 6.65 ± 0.41 Hz, n=25, P>0.05) and Cre^+^Fmr1^f/y^ (amplitude: 4.46 ± 0.19 pA; frequency: 6.68 ± 0.74 Hz, n=14, p>0.05) mice or between neurons from Cre^-^Fmr1^loxP-neo/y^ (amplitude: 4.41 ± 0.20 pA; frequency: 5.43 ± 0.36 Hz, n=25, P>0.05) and Cre^+^Fmr1^loxP-neo/y^ (amplitude: 4.4 ± 0.17 pA, n=16; frequency: 4.02 ± 0.61 Hz, n=16, p>0.05) (Fig. 1 D-F) mice. Thus, the enhanced NMDAR mEPSC amplitude observed in Fmr1 KO mice is unlikely to result from the selective loss of astrocytic FMRP.

**Figure 1.**
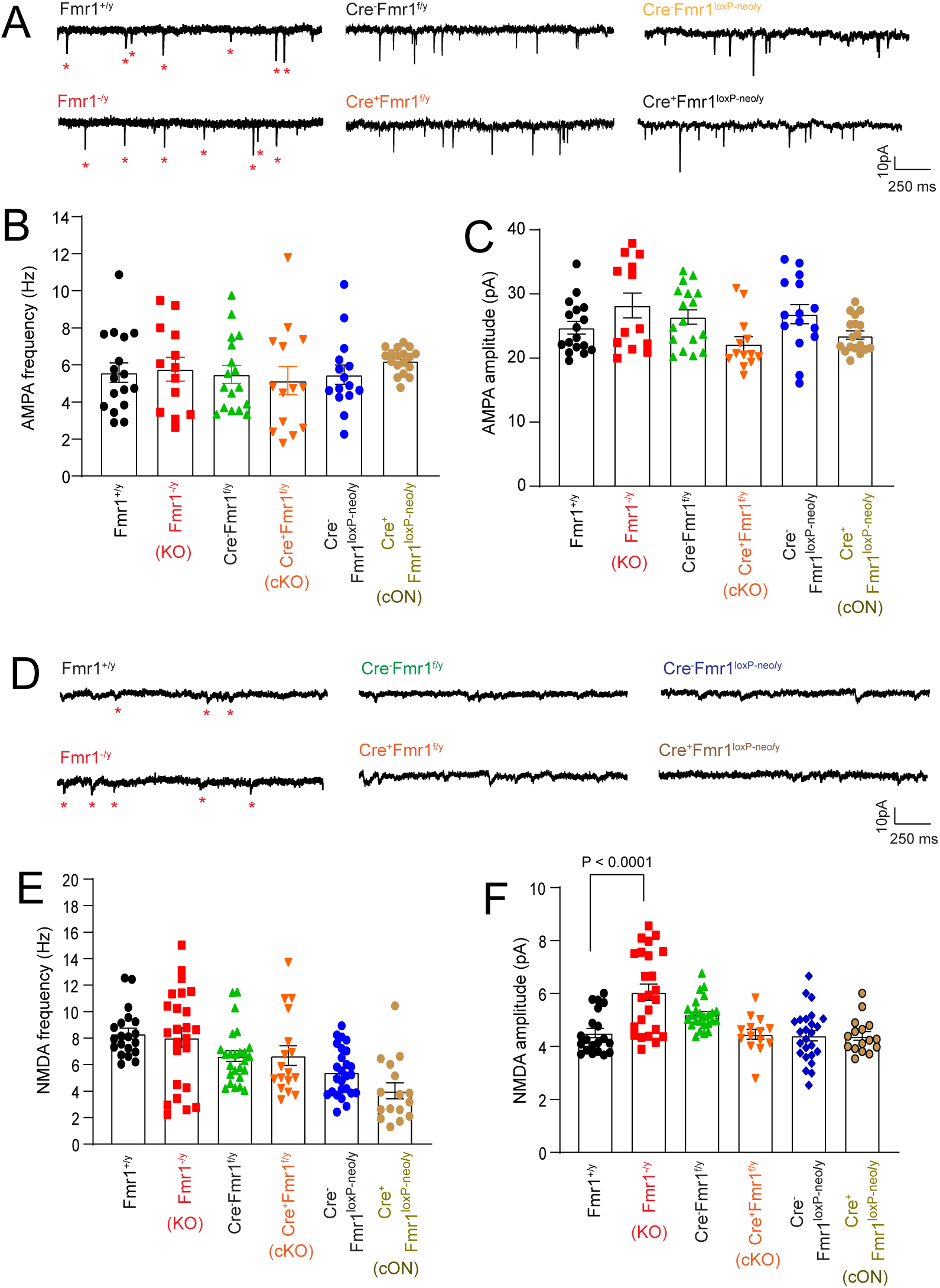
AMPAR and NMDAR-mediated spontaneous mEPSCs in cortical pyramidal neurons are not influenced by altered astroglial FMRP levels in astroglial Fmr1 cKO and cON mice. **A.** Representative AMPAR-mediated mEPSCs traces from pyramidal neurons of Fmr1 KO, astroglial Fmr1 cKO and cON cortical slices. Asterisks: individual AMPAR mEPSC; The average frequency (**B**) and amplitude (**C**) of AMPAR-mediated mEPSCs from pyramidal neurons of Fmr1 KO, astroglial Fmr1 cKO and cON cortical slices. N=13-18 neurons from 4-5 mice per group. P values were determined using one-way ANOVA with post-hoc Tukey’s test; **D**. Representative NMDAR-mediated mEPSCs traces from pyramidal neurons of Fmr1 KO, astroglial Fmr1 cKO and cON cortical slices. Asterisks: individual NMDAR mEPSC; The average frequency (**E**) and amplitude (**F**) of NMDAR-mediated mEPSCs from pyramidal neurons of Fmr1 KO, astroglial Fmr1 cKO and cON cortical slices. N= 16-25 neurons from 3-4 mice per group. P values were determined using one-way ANOVA with post hoc Tukey’s test.

As NMDAR-mediated mEPSC amplitude is typically small in Mg^2+^-free recordings (29), we also examined evoked NMDAR-mediated EPSCs of pyramidal neurons by using extracellular stimulation on somatosensory cortical slices from Fmr1 KO and astroglial Fmr1 cKO and cON mice. The AMPAR- and NMDAR-mediated responses were recorded at −70 mV or +40 mV respectively, as shown in representative traces (Fig. 2A-B). Our recordings showed that NMDAR-mediated responses were significantly larger (P<0.0001) in neurons from Cre^+^Fmr1^f/y^ mice (1.56 ± 0.02 pA/pF, n=33) in comparison to littermate control Cre^-^Fmr1^f/y^ mice (1.40 ± 0.02 pA/pF, n=34) (Fig. S1D), while AMPAR-mediated responses were highly comparable in these mice (Cre^+^Fmr1^f/y^, 2.40 ± 0.03 pA/pF, n=33; Cre^-^Fmr1^f/y^, 2.33 ± 0.03 pA/pF, n=34, p=0.1) (Fig. S1E). Similarly, the pairwise NMDAR- and AMPAR responses plot of individual neurons recorded exhibited a clear separation of NMDAR-but not AMPAR responses between Cre^+^Fmr1^f/y^ and Cre^-^ Fmr1^f/y^ cortical neurons (Fig. 2C). Importantly, NMDAR responses from Cre^+^Fmr1^f/y^ cortical neurons are highly comparable to those in Fmr1^-/y^ cortical neurons (Fig. 2D), implicating an important role of the loss of astroglial FMRP in enhancing evoked NMDA responses in Fmr1 KO mice. We then calculated the AMPA/NMDA current ratio and found that the AMPA/NMDA ratio of recorded cortical neurons in Cre^+^Fmr1^f/y^ mice (1.54 ± 0.02, n=33) is significantly lower (p<0.0001) than that in Cre^-^Fmr1^f/y^ cortical neurons (1.67 ± 0.01, n=34) (Fig. 2E). The application of dihydrokainate (DHK, 50 μM), a selective astroglial glutamate transporter GLT1 antagonist, on cortical slices further significantly (p<0.0001) decreased the AMPA/NMDA ratio in both Cre^+^Fmr1^f/y^ (1.36 ± 0.02, n=28) and Cre^-^Fmr1^f/y^ (1.52 ± 0.01, n=26) cortical neurons (Fig. 2E), indicating the involvement of an extracellular glutamate clearance mechanism in decreasing the AMPA/NMDA ratio. A similar reduction of the AMPA/NMDA ratio was also observed in Fmr1^-/y^ cortical neurons compared to Fmr1^+/y^ cortical neurons in untreated (Fmr1^+/y^, 1.61 ± 0.02, n=24; Fmr1^-/y^, 1.56 ± 0.01, n=24, p=0.01) and DHK-treated (Fmr1^+/y^, 1.55 ± 0.01, n=20; Fmr1^-/y^, 1.48 ± 0.01, n=20, p=0.0003) slices (Fig. 2F). These results further suggest that selective loss of astroglial FMRP alone is sufficient to fully recapitulate the decreased synaptic AMPA/NMDA ratio observed in Fmr1 KO mice. Indeed, evoked NMDAR- (Cre^-^Fmr1^loxP-neo/y^, 1.60 ± 0.02 pA/pF, n=32; Cre^+^Fmr1^loxP-neo/y^, 1.48 ± 0.02 pA/pF, n=33, p<0.0001) (Fig. S1F) but not AMPAR- (Cre^-^ Fmr1^loxP-neo/y^, 2.47 ± 0.03 pA/pF, n=32; Cre^+^Fmr1^loxP-neo/y^, 2.41 ± 0.03 pA/pF, n=33, p=0.19) responses in Cre^+^Fmr1^loxP-neo/y^ cortical neurons are significantly lower than that in control Cre^-^ Fmr1^loxP-neo/y^ cortical neurons (Fig. S1G). A clear downward shift (decrease) of NMDA amplitude in individually recorded Cre^+^Fmr1^loxP-neo/y^ cortical neurons compared to Cre^-^Fmr1^loxP-neo/y^ cortical neurons was also observed (Fig. 2G). Consequently, the lower AMPA/NMDA ratio observed in control Cre^-^Fmr1^loxP-neo/y^ mice (1.54 ± 0.02, n=33) with minimal FMRP levels (26), which is also comparable with that in Fmr1^-/y^ mice (1.56 ±0.01, n=24), is significantly (p=0.004) increased in Cre^+^Fmr1^loxP-neo/y^ mice (1.67 ± 0.01, n=34) in which FMRP is selectively re-expressed in astroglia (Fig. 2H). The DHK treatment, while reducing the overall AMPA/NMDA ratio in both Cre^-^ Fmr1^loxP-neo/y^ and Cre^+^Fmr1^loxP-neo/y^ mice when compared to the untreated condition, still shows a significant (p<0.0001) increase of the AMPA/NMDA ratio in Cre^+^Fmr1^loxP-neo/y^ cortical neurons (1.52 ± 0.01, n=26) in comparison to Cre^-^Fmr1^loxP-neo/y^ cortical neurons (1.36 ± 0.02, n=28) (Fig. 2H). Interestingly, the decrease of the AMPA/NMDA ratio appears to be age-dependent, as the AMPA/NMDA ratio shows the decreasing trend but is not statistically different (p=0.06) between Cre^+^Fmr1^f/y^ (1.63 ± 0.03, n=10) and Cre^-^Fmr1^f/y^ (1.70 ± 0.02, n=10) cortical neurons at younger ages (P24-30) (Fig. S1H).

**Figure 2.**
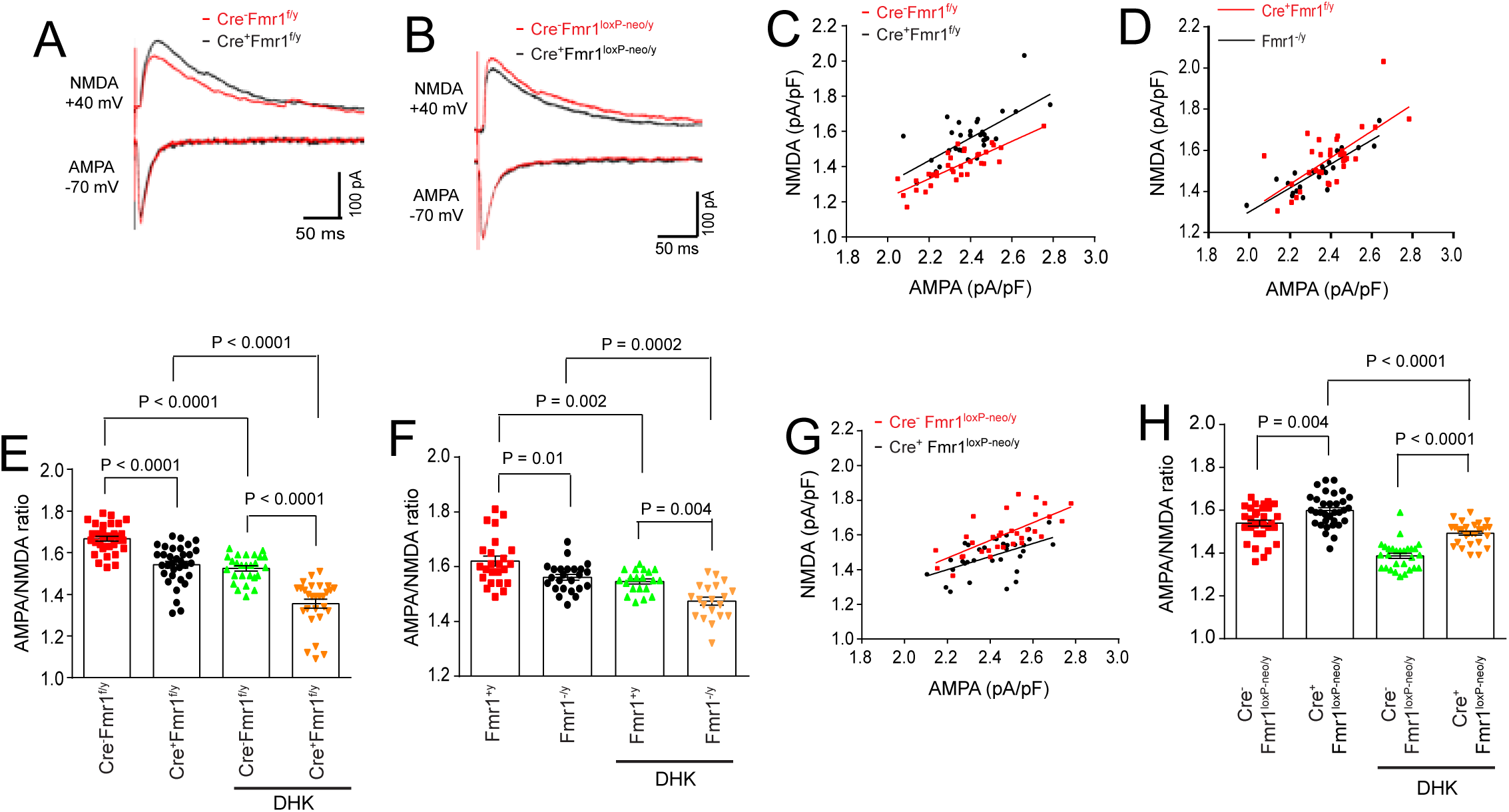
Modulation of evoked AMPA/NMDA ratio of cortical neurons by astroglial FMRP in astroglial Fmr1 cKO and cON mice. Representative evoked AMPAR-(−70 mV) and NMDAR-(+40 mV) mediated EPSC traces from astroglial cKO (**A**, Cre^+^Fmr1^f/y^ and Cre^-^Fmr1^f/y^) and cON (**B**, Cre^+^Fmr1^loxP-neo/^y and Cre^-^Fmr1^loxP-neo/y^) mice. **C.** Scatter plot of individual NMDAR and AMPAR responses from patched cortical pyramidal neurons in Cre^-^Fmr1^f/y^ (red) and Cre^+^Fmr1^f/y^ (black) mice. N=33-34 neurons from 6-8 mice per group; **D.** Scatter plot of individual NMDAR and AMPAR responses from patched cortical pyramidal neurons in Cre^+^Fmr1^f/y^ (red) and Fmr1^-/y^ mice (black). N=24-33 neurons from 6-8 mice per group. No difference was observed between Cre^+^Fmr1^f/y^ and Fmr1^-/y^ mice. **E.** The average AMPA/NMDA ratio of cortical neurons in untreated and 50 µM DHK-treated slices from Cre^-^Fmr1^f/y^ and Cre^+^Fmr1^f/y^ mice. N=26-34 neurons from 6-8 mice per group, P values were determined using one-way ANOVA with post hoc Tukey’s test. **F.** The average AMPA/NMDA ratio of cortical neurons in untreated and 50 µM DHK-treated slices from Fmr1^+/y^ and Fmr1^-/y^ mice. N=20-24 neurons from 6-8 mice per group, P values were determined using one-way ANOVA with post hoc Tukey’s test. **G.** Scatter plot of NMDAR and AMPAR responses from patched cortical pyramidal neurons in Cre^-^Fmr1^loxP-neo/y^ (red) and Cre^+^Fmr1^loxP-neo/y^ (black) mice. N=32-33 neurons from 6-8 mice per group, **H.** The average AMPA/NMDA ratio of cortical neurons in untreated and 50 µM DHK-treated slices from Cre^-^ Fmr1^loxP-neo/y^ and Cre^+^Fmr1^loxP-neo/y^ mice. N=28-33 neurons from 6-8 mice per group, P values were determined using one-way ANOVA with post hoc Tukey’s test.

### Elongated spontaneously occurring cortical UP states in astroglial Fmr1 cKO mice

Circuit level hyperexcitability is considered one of the central mechanisms underlying typical FXS symptoms such as hyperactivity, seizures, and sensory hypersensitivity (30). Cortical circuit hyperexcitability has been observed in Fmr1 KO mice *in vitro* and *in vivo* (31–33). In particular, the duration of the firing phase of the UP state becomes prolonged in somatosensory cortical neurons of Fmr1 KO mice (32). The UP state often refers to a short period of synchronized neuronal network activity (action potential firing) that is formed as a result of the net balance of excitation and inhibition of cortical circuits (34). Although the selective deletion of the *fmr1* gene in cortical excitatory neurons using the Emx1-Cre driver line was sufficient to cause prolonged UP states (32), the Emx1 gene is equivalently expressed in both neurons and astroglia based on RNA-seq data of brain cells (35), making it possible that the loss of astroglial FMRP may contribute to prolonged UP states. Additionally, stimulating a single astroglia activates other astroglia in the local circuit and is sufficient to trigger UP state synchronization of neighboring neurons (36). Here, we measured UP states with extracellular recording in layer 4 of somatosensory cortex from Fmr1 KO, astroglial Fmr1 cKO, and cON mice and their respective littermate control mice. UP states were defined as previously reported (32) and examples of expanded traces of UP states are shown in Fig. S2A. We observed that the duration of spontaneously occurring UP states was significantly elongated (p=0.0002) in Fmr1^-/y^ (788.1 ± 78.6 ms, n=10) compared to in Fmr1^+/y^ mice (382.1 ± 31.6 ms, n=8) (Fig. 3A-B). The upward shape of UP states in our recordings, as compared to the biphasic shape of UP states in previous studies (32, 37, 38), is primarily due to the difference in the resistance (tip size) of the recording electrode used (∼1 MΩ in our recordings vs. ∼0.5 MΩ in previous recordings (37, 38)), as shown in Fig. S2B. The difference in the electrode resistance has no effect on the duration and frequency of UP states (Fig. S2C-D). A similarly elongated duration of UP states was also observed in astroglial Fmr1 cKO (Cre^+^Fmr1^f/y^) mice in comparison with their littermate control Cre^-^Fmr1^f/y^ mice (Cre^-^Fmr1^f/y^: 407.8 ± 16.7 ms, n=21; Cre^+^Fmr1^f/y^: 721.2 ± 78.0 ms, n=15, p<0.0001) (Fig. 3C-D). Notably, the increased duration of UP states is fully attenuated in astroglial Fmr1 cON (Cre^+^Fmr1^loxP-neo/y^) mice when compared to their control Cre^-^Fmr1^loxp-neo/y^ mice (Cre^-^Fmr1^loxP-neo/y^: 611.1 ± 28.8 ms, n=18; Cre^+^Fmr1^loxP-neo/y^: 395.1± 17.2 ms, n=16, p<0.0001) (Fig. 3E-F). In contrast to the duration changes, we found no significant changes in the frequency of UP states from all recordings (Fmr1^+/y^: 0.1 ± 0.015 Hz, n=8; Fmr1^-/y^: 0.08 ± 0.01, n=10; Cre^-^Fmr1^f/y^: 0.08 ± 0.004 Hz, n=21; Cre^+^Fmr1^f/y^: 0.08 ± 0.004, n=15; Cre^-^Fmr1^loxP-neo/y^: 0.07 ± 0.003 Hz, n=18; Cre^+^Fmr1^loxP-neo/y^: 0.08 ± 0.005, n=16) which is consistent with other recordings from layer 4/5 neurons in Fmr1 KO mice (32). These results show that the selective loss of FMRP in astroglia is sufficient to induce significantly longer UP states, consistent with the enhanced evoked NMDA responses in Fig. 2. These results indicate potential astroglia-mediated mechanisms that contribute to the cortical circuit hyperexcitability observed in Fmr1 KO mice.

**Figure 3.**
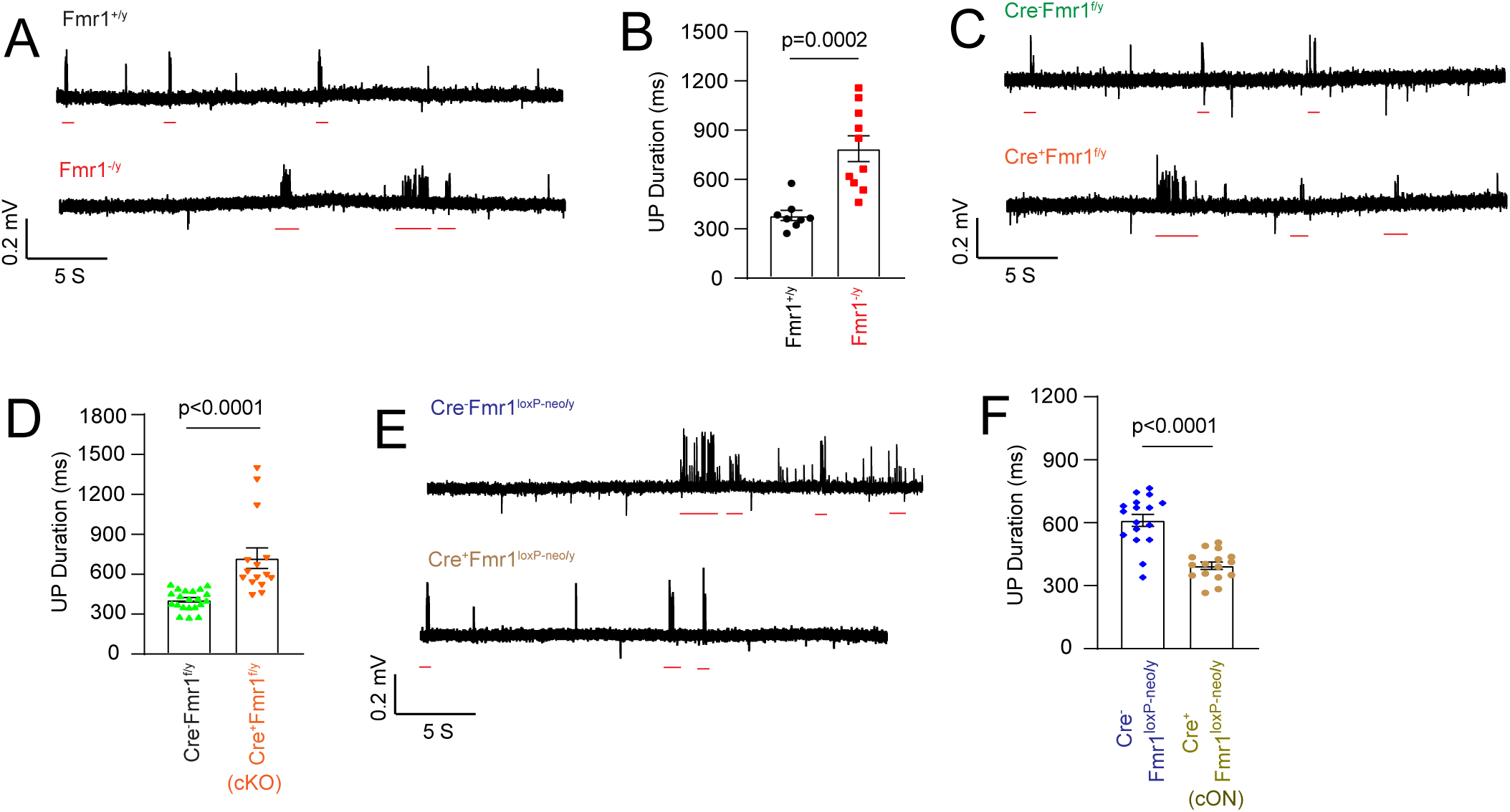
Modulation of cortical UP state duration by astroglial FMRP in astroglial Fmr1 cKO and cON mice. Representative recordings (**A**) and average duration (**B**) of spontaneous UP states from Fmr1^+/y^ and Fmr1^-/y^ mice. N=8-10 slices from 4 mice per group. Representative recordings (**C**) and average duration (**D**) of spontaneous UP states from Cre^+^Fmr1^f/y^ and Cre^-^Fmr1^f/y^ mice. N=15-21 slices from 6 mice per group. Representative recordings (**E**) and average duration (**F**) of spontaneous UP states from Cre^-^Fmr1^loxP-neo/y^ and Cre^+^Fmr1^loxP-neo/y^ mice. N=16-18 slices from 6 mice per group. ms: millisecond; P values were determined using the two tailed unpaired t-test.

### Selective loss of astroglial FMRP impacts typical FXS-relevant behavioral and learning phenotypes

FXS is commonly manifested with several behavioral phenotypes such as hyperactivity and social avoidance, that have been recapitulated in Fmr1 KO mice in the open field and three-chamber social-interaction tests, respectively (39, 40). It remains unexplored how the loss of FMRP in different brain regions/cell types impacts these behavioral phenotypes. As cortex is one of the primary brain regions involved in these tests and cortical synaptic/circuitry activity is altered in astroglial Fmr1 cKO mice (Figs. 2-3), we decided to test whether selective loss of astrocytic FMRP influences these cortical circuit-centered behaviors. We first tested locomotor activity in the open field test by analyzing the total running distance from the first 10 minutes in the open field arena. Because mice tend to explore the new environment, the first 10 minutes of the open field test typically best reflect the locomotor activity of individual mice, as shown in the representative running heatmaps (Fig. 4A). Although there is a clear trend for increased overall distance traveled by Cre^+^Fmr1^f/y^ (3634 ± 350 cm, n=12) mice in comparison to Cre^-^Fmr1^f/y^ (3035 ± 369 cm, n=13) mice, the difference in the total travel distance is not statistically significant (p=0.26). In contrast, the overall distance traveled by Cre^+^Fmr1^loxP-neo/y^ (3162 ± 179 cm, n=16) mice is significantly (p=0.02) decreased compared to Cre^-^Fmr1^loxP-neo/y^ (4474 ± 248 cm, n=9) mice. Subsequent analysis of the running distance in each 2-minute block showed that the distance traveled by Cre^+^Fmr1^f/y^ mice is consistently longer, though not significant except for the last 2-minutes block (P=0.01), than that traveled by Cre^-^Fmr1^f/y^ mice at the same time block, confirming an overall tendency of elevated locomotor activity of Cre^+^Fmr1^f/y^ mice (Fig. 4C). Importantly, similar 2 minutes block analysis in astroglial Fmr1 cON mice showed that the re-expression of FMRP in astroglia consistently reduces the running distances in early time blocks by Cre^+^Fmr1^loxP-neo/y^ mice in comparison to Cre^-^Fmr1^loxP-neo/y^ mice in which minimal FMRP is expressed (Fig. 4D).

**Figure 4.**
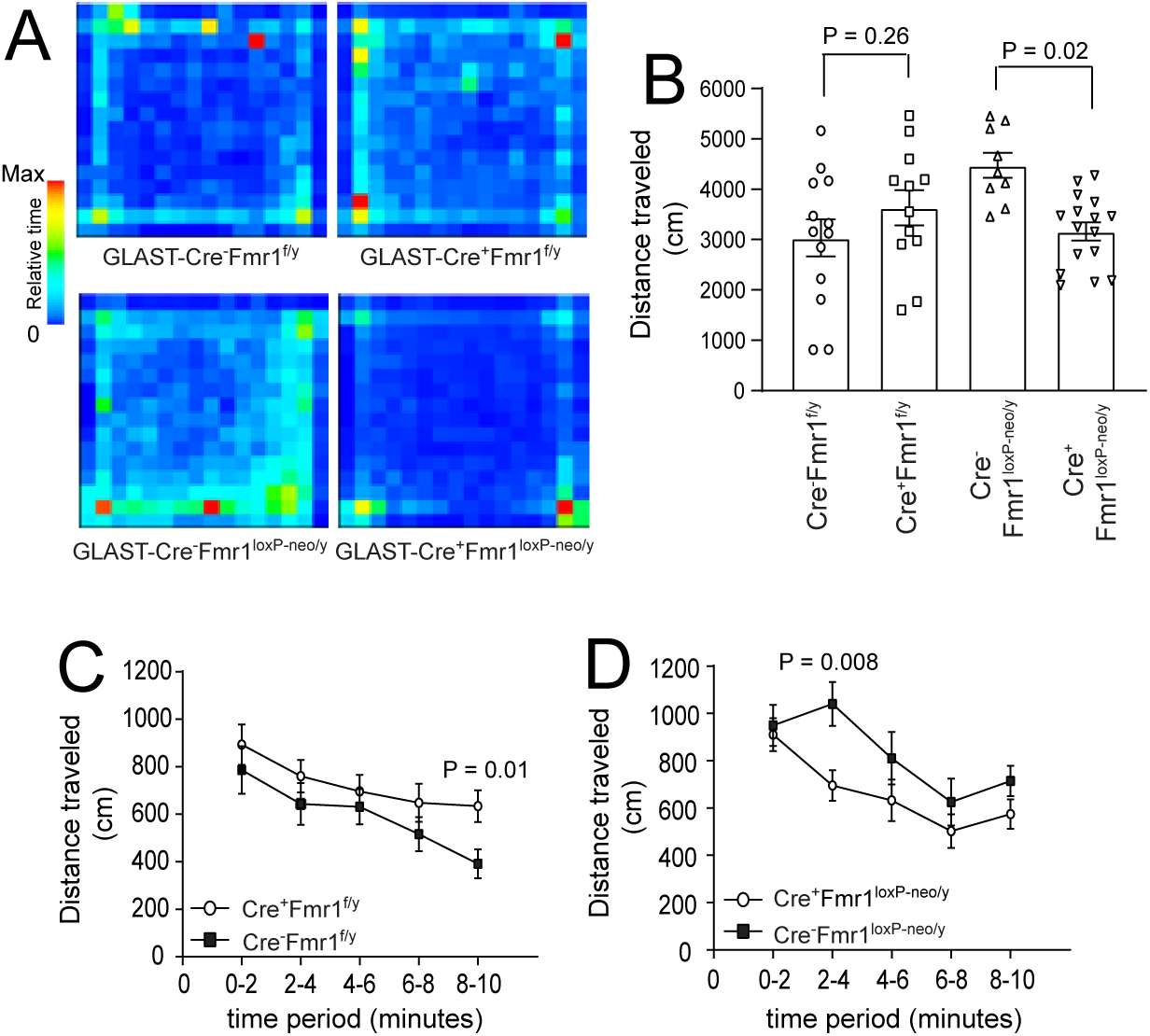
Re-expression of FMRP sufficiently reduces locomotor hyperactivity in astroglial Fmr1 cON mice. **A.** Representative heat maps showing time spent at each position of the open field box from astroglial cKO (Cre^-^Fmr1^f/y^ and Cre^+^Fmr1^f/y^) and cON (Cre^-^Fmr1^loxP-neo/y^ and Cre^+^Fmr1^loxP-neo/y^) mice. **B.** Total distance traveled by astroglial Fmr1 cKO and cON mice during the first 10 minutes in the open field box. N=9-16 mice per group, P values were determined by two-tailed unpaired t-test. Distance traveled in each 2-minute interval of the open field test in astroglial cKO (**C**) and cON (**D**) mice. N=9-16 mice per group, P values were determined using two tailed unpaired t-test.

We next performed the three-chamber social interaction test with astroglial Fmr1 cKO and cON mice to assess the influence of astroglial FMRP on cognition in the aspect of general sociability (interest) and social novelty (novel vs. familiar). As shown in Fig. 5A, Cre^+^Fmr1^f/y^ mice (empty chamber: 196.6 ± 24.7 sec, n=17; stranger mouse: 468.8 ± 29.4 sec, n=17, p<0.0001) showed normal sociability as their control Cre^-^Fmr1^f/y^ mice (empty chamber: 177.4 ± 6.9 sec, n=14; stranger mouse: 446.6 ± 28.2 sec, n=12, p<0.0001) with a clear and strong preference for interacting with the stranger mouse over the empty chamber. A similar preference for the stranger mouse over the empty chamber was also observed with Cre^-^Fmr1^loxP-neo/y^ (empty chamber: 221.6±24.4 sec, n=11; stranger mouse: 428.8±26.2 sec, n=11, p=0.004) and Cre^+^Fmr1^loxP-neo/y^ mice (empty chamber: 264.1 ±40.4 sec, n=15; stranger mouse: 490.7 ± 43.7, n=15, p=0.0002) (Fig. 5B). Interestingly, although the littermate control Cre^-^Fmr1^f/y^ mice have a clear preference (p=0.005) for the novel (316.6 ± 21.8 sec, n=14) stranger mouse over the familiar (224.5 ± 20.4 sec, n=14) stranger mouse (Fig. 5C), a natural preference in rodents, Cre^+^Fmr1^f/y^ mice showed no clear preference (p=0.08) for either the novel (323 ± 22.56 sec, n=17) or the familiar (268 ± 20.44 sec, n=17) stranger mouse (Fig. 5C). The re-expression of FMRP in astroglia in Cre^+^Fmr1^loxP-neo/y^ mice, on the other hand, sufficiently restores the preference (p=0.006) for the novel (407.7 ±38.8 sec, n=14) stranger mouse over the familiar (269.9 ± 24.7 sec, n=14) stranger mouse (Fig. 5D), while the control Cre^-^Fmr1^loxP-neo/y^ mice (with minimal FMRP levels) have no clear preference (p=0.09) for either the novel (347.4 ± 43.4 sec, n=13) or the familiar (249.4 ± 33.2 sec, n=13) stranger mouse (Fig. 5D). These results demonstrate that the absence of FMRP in astroglia contributes to the social novelty deficit of Fmr1 KO mice.

**Figure 5.**
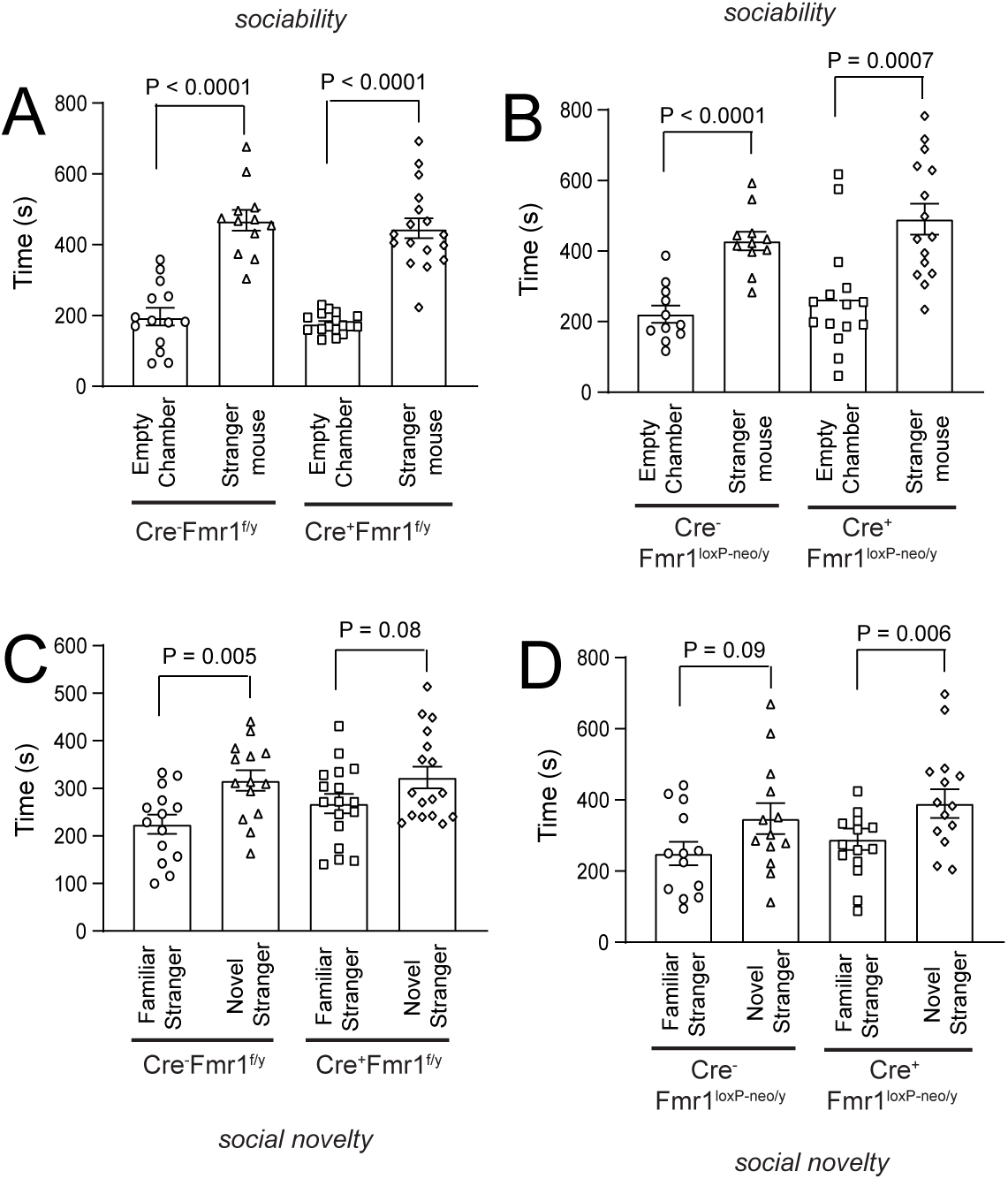
Astroglial FMRP significantly contributes to social novelty deficits in astroglial Fmr1 cKO mice. Sociability test (stranger mouse vs. empty chamber) of astroglial cKO (**A,** Cre^-^Fmr1^f/y^) and cON (**B,** Cre^-^Fmr1^loxP-neo/y^) mice with their respective controls (Cre^+^Fmr1^f/y^ or Cre^+^Fmr1^loxP-neo/y^). N=11-17 mice per group, P values determined using two-tailed unpaired t-test. Social novelty (familiar vs. novel mouse preference) test of astroglial cKO (**C,** Cre^-^Fmr1^f/y^) and cON (**D,** Cre^-^ Fmr1^loxP-neo/y^) mice with their respective controls (Cre^+^Fmr1^f/y^ or Cre^+^Fmr1^loxP-neo/y^). N=13-17 mice per group, P values determined using two-tailed unpaired t-test. s: seconds;

In addition to the hyperactivity and social avoidance, learning disability and impaired memory is another significant feature in FXS. Previous studies have shown that Fmr1 KO mice exhibit impaired cognitive functions in a variety of cognitive tests including Barnes maze, novel task discrimination, and inhibitory avoidance (IA) (41, 42). Whether the loss of astroglial FMRP contributes to impaired cognitive functions in Fmr1 KO mice (or FXS) remains unexplored. The inhibitory avoidance and extinction test involve both memory acquisition and extinction processes (Fig. 6A) that allow dissection of specific deficits to which the selective loss of astroglial FMRP may contribute. We therefore decided to perform the inhibitory avoidance test on astroglial Fmr1 cKO and cON mice. As shown in Fig. 6B, both Cre^-^Fmr1^f/y^ (6h: 279.2 ± 51.8 sec vs. 0h: 24.7 ± 3.9 sec) and Cre^+^Fmr1^f/y^ (6h: 151.4 ± 34.9 sec vs. 0h: 36 ± 7.4 sec) mice are able to acquire memory of the aversive shock by showing significantly increased latency to enter the dark chamber at 6h post-shock. However, Cre^+^Fmr1^f/y^ mice (6h: 151.4 ± 34.9 sec) have a significantly shorter latency than that of control Cre^-^Fmr1^f/y^ mice (6h: 279.2 ± 51.8 sec, Fig. 6B). In addition, majority (60%) of Cre^+^Fmr1^f/y^ (Fig. 6C) mice, but only 30% of control Cre^-^Fmr1^f/y^ mice (Fig. 6D), entered the dark chamber within 100sec at the 6h following the shock. Moreover, only 6.7% (1/15 mice, Fig. 6E) of Cre^-^Fmr1^f/y^ mice did not acquire the contextual fear memory (no significant increase of latency time at 6h vs. 0h) following the initial shock while 33.3% (7/21, Fig. 6F) of Cre^+^Fmr1^f/y^ mice did not acquire the contextual fear memory. These results indicate a deficit in memory acquisition in the absence of astroglial FMRP in Cre^+^Fmr1^f/y^ mice. The latency to enter the dark chamber in Cre^-^Fmr1^f/y^ mice (24h: 261.3 ± 63.6 sec; 48h: 194.4 ± 55.5 sec) is consistently and significantly (p=0.01 and p=0.03 respectively) longer than that in Cre^+^Fmr1^f/y^ mice (24h: 105.1 ± 27.1 sec; 48h: 81.1 ± 20 sec) during the memory extinction phase (24h and 48h, Fig. 6B), suggesting that the memory extinction also becomes exaggerated (or less memory retention) in Cre^+^Fmr1^f/y^ mice, as previously reported in Fmr1 KO mice (42). Latency changes from 6h to 24h of individual Cre^-^Fmr1^f/y^ and Cre^+^Fmr1^f/y^ mice tested were shown in Fig. S3A-B. We next performed inhibitory avoidance test on astroglial cON mice and found that both Cre^-^Fmr1^loxP-neo/y^ (6h: 141.6± 41.9 sec; 24h: 24h: 59.9 ± 22.9 sec; 48h: 43.4 ± 18.1 sec) and Cre^+^Fmr1^loxP-neo/y^ (6h: 136.2 ± 29.9 sec; 24h: 79.2 ± 18.3 sec; 48h: 61.5 ± 15.8 sec) mice showed no significant difference in latency time at 6h, 24h, and 48h after the initial shock (Fig. 6G), though both Cre^-^Fmr1^loxP-neo/y^ and Cre^+^Fmr1^loxP-neo/y^ mice are able to acquire the IA memory (Fig. 6G). The cumulative probability of latency from Cre^-^Fmr1^loxP-neo/y^ and Cre^+^Fmr1^loxP-neo/y^ mice at both 6h (Fig. 6H) and 24h (Fig. 6I) is also similar. Latency changes from 6h to 24h of individual Cre^-^Fmr1^loxP-neo/y^ and Cre^+^Fmr1^loxP-neo/y^ mice tested were shown in Fig. S3C-D. The number of Cre^-^Fmr1^loxP-neo/y^ (13.3%, 2/15, Fig. 6J) and Cre^+^Fmr1^loxP-neo/y^ (18.2%, 4/22, Fig. 6K) mice that did not learn the avoidance behavior following the initial shock is also comparable.

**Figure 6.**
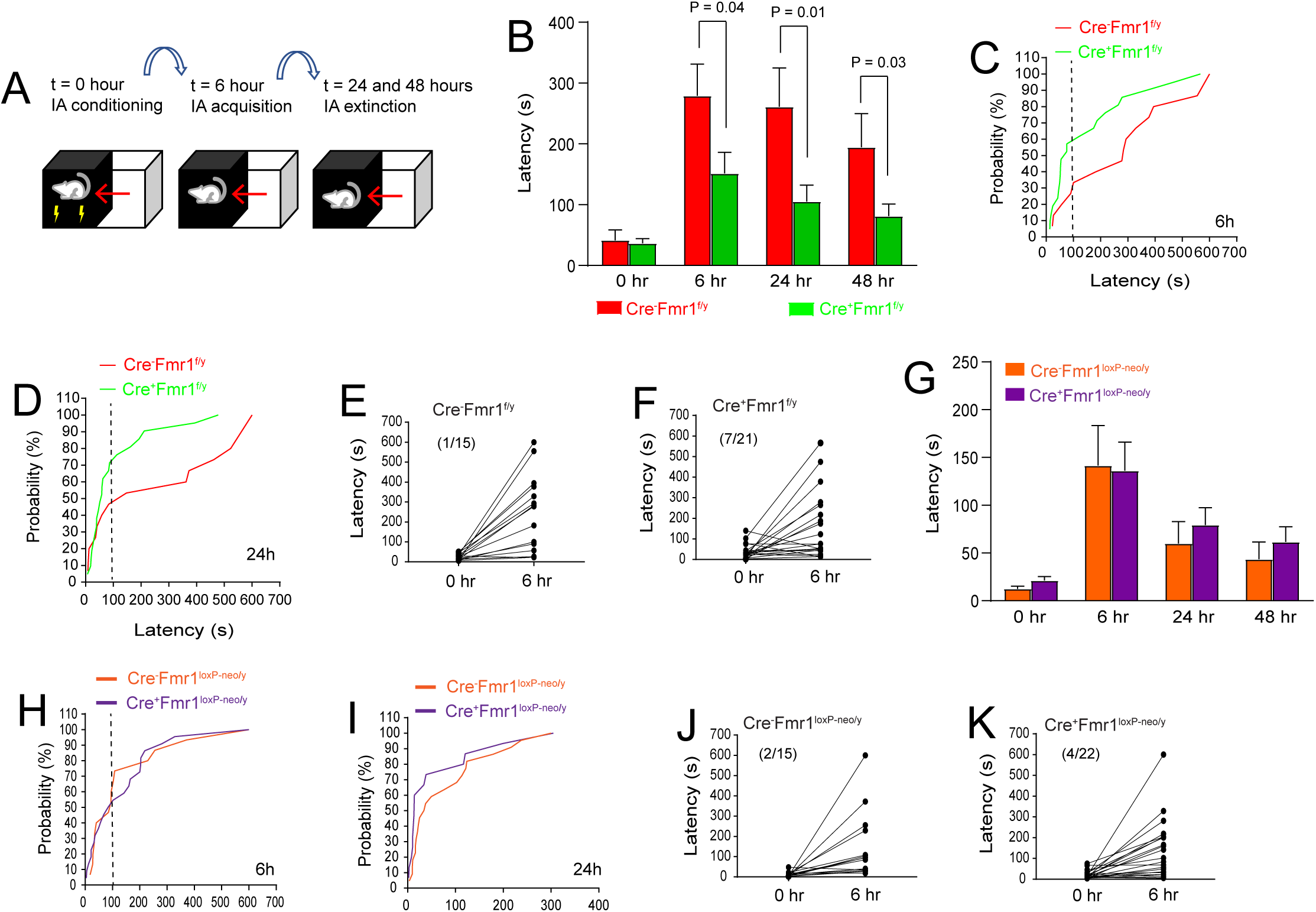
Selective loss of astroglial Fmr1 induces deficits in memory acquisition and extinction during inhibitory avoidance extinction task. **A.** Schematic diagram for the IA test. Mice were given the IA training at time 0 and the latency to enter the dark side was measured at 6 h (post-acquisition). They were then given IA extinction training, and latency was again measured at 24h. Testing was followed by another round of IA extinction training, and latency was measured again at 48 h. **B.** Average latency to enter the dark chamber between Cre^-^Fmr1^f/y^ and Cre^+^Fmr1^f/y^ mice at 6 h, 24 h, and 48 h. N=15-21 mice per group, P values determined using multiple unpaired two-tailed t-tests. s: seconds; Cumulative probability curve of the latency of Cre^-^Fmr1^f/y^ and Cre^+^Fmr1^f/y^ mice at 6h (**C**) or 24h (**D**) post IA training (foot shock). N=15-21 mice per group, Raw data of latency changes from 0h to 6h in Cre^-^Fmr1^f/y^ (**E**) and Cre^+^Fmr1^f/y^ (**F**) mice. Each line indicates a single mouse. **G.** Average latency to enter the dark chamber between Cre^-^Fmr1^loxP-neo/y^ and Cre^+^Fmr1^loxP-neo/y^ mice at 6h, 24h, and 48h. N=15-22 mice per group, P values determined using multiple unpaired two-tailed t-test. Cumulative probability curve of the latency of Cre^-^Fmr1^loxP-neo/y^ and Cre^+^Fmr1^loxP-neo/y^ at 6h (**H**) or 24h (**I**) post IA training (foot shock). N=15-22 mice per group, Raw data of latency changes from 0h to 6h in Cre^-^Fmr1^loxP-neo/y^ (**J**) and Cre^+^Fmr1^loxP-neo/y^ (**K**) mice. Each line indicates a single mouse.

## Discussion

Our current studies show that selective loss of astroglial FMRP contributes to cortical hyperexcitability by enhancing NMDAR-mediated evoked but not spontaneous mEPSCs and by elongating cortical UP state duration. Our results also show that selective loss of astroglial FMRP is sufficient to increase (though not statistically significant) locomotor activity, significantly diminish social novelty preference, and induce memory acquisition and extinction deficits in astroglial Fmr1 cKO mice. Importantly, re-expression of astroglial FMRP is able to rescue the hyperactivity and social novelty phenotypes in astroglial Fmr1 cON mice. These results, achieved by selectively modulating FMRP expression levels in astroglia, fill the missing gap as to whether non-neuronal (astro)glial cells are involved in modulating FXS relevant synaptic signaling and behavior phenotypes. These results also present compelling evidence in considering astroglia for testing the FMRP reactivation strategy in the future.

Although NMDAR responses of cortical pyramidal neurons in both evoked and spontaneous conditions were enhanced in Fmr1 KO mice in our results, enhanced NMDAR response was only observed in the evoked (but not spontaneous) condition in astroglial Fmr1 cKO mice that was fully attenuated in astroglial Fmr1 cON mice. This discrepancy in NMDA responses between Fmr1 KO and astroglial Fmr1 cKO mice suggests that the spontaneous NMDAR mEPSC response is likely to be primarily regulated by FMRP in neurons while astroglial FMRP only contributes to evoked NMDA responses. Extracellular stimulation during the evoke procedure typically elicits a large set of synapses with much more robust release of glutamate and potential glutamate spillover, adding a significant burden for its timely and proper clearance. We previously showed that functional expression of astroglial GLT1, one of the dominant glutamate transporters responsible for clearing extracellular glutamate, is dysregulated in Fmr1 KO and astroglial Fmr1 cKO mice (20, 43). Thus, the compromised glutamate handling (due to GLT1 dysregulation) may not be able to efficiently and quickly clear elevated extracellular glutamate levels from evoked stimulation, which leads to enhancement of evoked (but not spontaneous) NMDAR responses in Fmr1 KO and astroglial Fmr1 cKO mice. Like evoked stimulation, rhythmic neuronal network activity also triggers the release of large amount of glutamate during UP states. Thus, GLT1 dysregulation in FXS models may similarly lead to improper and, specifically, delayed clearance of large amount of synaptically released glutamate during UP states, contributing to the elongated firing phase during the UP state. Another interesting study found that astroglial activation can trigger UP state synchronization of neighboring neurons, in which pharmacological inhibition of GLT1 also led to increased UP states (36). As UP states resemble cortical rhythmic states during relaxed behavioral states and sleep (44), our current study and others may shed a new light on how astroglia are involved in modulating these states/behaviors. Interestingly, selective deletion of FMRP in cortical interneurons has no obvious effect on UP state duration (32). Although the selective deletion of FMRP in excitatory neurons has been shown to be sufficient to elongate UP state duration using the Emx1-Cre driver line (32), a number of cell types including radial glia, Cajal-Retzius cells, glutamatergic neurons, astroglia, and oligodendrocytes can be derived from Emx1-expressing linage (45), casting considerable doubt on the involvement of excitatory neuronal FMRP in elongating UP state duration.

Our use of Bac *slc1a3*-CreER transgenic mice and the 4-OHT administration paradigm (P4-P9 injections) for generating astroglial Fmr1 cKO and cON mice leads to highly specific Fmr1 deletion in astroglia as radial glial cells in which the *slc1a3* promoter is also active are all destined to differentiate into astroglia during the first postnatal week (46). We have also previously confirmed the specificity of *slc1a3*-CreER-mediated recombination in astroglia (20). Although a moderate recombination efficiency of a knock-in *slc1a3*-CreER driver line (47) was found in adult hippocampal astroglia based on GCaMP6 induction (48), the *slc1a3* promoter is mostly active during postnatal development with a > 80% recombination efficiency in astroglia based on tdTomato reporter expression in Ai14 reporter mice (20, 49). The single copy of the loxP floxed *fmr1* or *neo* gene on the X chromosome in Fmr1^f/y^ or Fmr1^loxP-neo/y^ mice, respectively, also facilitates the efficient diminishment or restoration of FMRP expression in astroglial Fmr1 cKO or cON mice, as shown in the FMRP immunoblot from FAC sorted cortical astroglia in astroglial Fmr1 cKO or cON mice. In addition, we are able to reversibly induce and attenuate NMDAR synaptic signaling and several FXS-related behaviors examined (except IA memory acquisition or extinction) in astroglial Fmr1 cKO and cON mice, respectively. As the IA training is a primarily hippocampus-centered task, the relevant hippocampal circuitry may also be differentially influenced by the selective deletion or restoration of astroglial FMRP.

Astroglia have been increasingly demonstrated to modulate cognitive functions and animal behaviors, such as emotion, motor activity and coordination, and sensory processing (50). Astrocytic glycogen breakdown and lactate shuttle to neurons are essential for long-term memory formation (51). Complete deletion of the *aqp4* gene (encoding aquaporin 4), a gene highly and selectively expressed in astroglia, leads to reduced long-term memory that can be rescued by increasing GLT1 expression (52). Cerebellar astroglia (Bergmann glia) express AMPA receptors that are required for fine motor coordination (53), and astroglia-derived ATP mediates antidepressant-like effects in mouse models of depression (54). Our current study presents direct evidence that astroglial FMRP significantly modulates FXS-related behavioral and learning phenotypes, especially social memory and cognitive functions. Although how astroglia modulate FXS-related behaviors remains unknown, substantial progress has been made in defining astroglia-mediated mechanisms in modulating synaptic connectivity and plasticity in the developing and adult CNS (55). These mechanisms provided high value targets/pathways to mechanistically understand the pathophysiological involvement of astroglia in FXS-related cognitive impairments and behavior alterations. Indeed, alterations of astroglia-secreted synaptogenic signals have been observed in Fmr1 KO mice (56). We also previously showed dysregulated mGluR5 signaling in Fmr1 KO astroglia (43). In addition, profiling of FMRP bound mRNAs from brain tissue has identified a number of mRNAs that are highly and selectively enriched in astroglia (4), including *slc1a2* (encoding GLT1), *glul* (encoding glutamine synthetase), *aldoc* (encoding aldolase), *apoe* (encoding apolipoprotein E), *sparcl1* (encoding hevin), and others, suggesting a strong molecular involvement of astroglia in FXS pathogenesis. Future explorations to define the roles of these (and additional) important astroglial genes in FXS will provide mechanistic insights into the astroglial contribution to FXS and new opportunities for therapeutic interventions in FXS.

## Materials and Methods

### Animals

The Fmr1^f/f^ and Fmr1^loxP-neo/loxP-neo^ mice were generated as previously described (26). Bac *slc1a3* CreERT transgenic mice (C57 BL6 background, stock#: 012586), Fmr1 KO mice (FVB background, stock#: 003024) were obtained from the Jackson Laboratory. The Fmr1^f/f^ and Fmr1^loxP-neo/loxP-neo^ mice were bred with Bac *slc1a3* CreERT transgenic mice to generate inducible astroglia specific cKO and cON mice. We used only male mice from Fmr1 KO, astroglial Fmr1 cKO and cON genotypes in all experiments because the *fmr1* locus is on the X chromosome and males are therefore more severely affected than females among FXS patients and in mouse models. All mice were maintained on a 12h light/dark cycle with food and water ad libitum. Care and treatment of animals in all procedures strictly followed the NIH Guide for the Care and Use of Laboratory Animals and the Guidelines for the Use of Animals in Neuroscience Research and the Tufts University IACUC.

### Drug administration

Tamoxifen (4-OHT) (Sigma) was resuspended at 20 mg/ml in ethanol and diluted into sunflower seed oil at a final concentration of 2 mg/ml in 10% of ethanol. For astroglial Fmr1 cKO and cON mice, daily intraperitoneal injections (IP) of 20 μl 4-OHT (50 mg/kg) were administered from P4 to P9 for a total dose of 0.25 mg. All littermate control mice received the same 4-OHT injections as the experimental mice.

### Cortical brain slice preparation

Cortical brain slices were prepared from juvenile (P24-30) and young adult (P38-45) male WT and Fmr1 KO, astroglial Fmr1 cKO, and cON mice according to methods previously described (20). Briefly, animals were anesthetized with ketamine/xylazine cocktail (110 mg/kg/10mg/kg); cortex was quickly removed and 300 μm cortical slices were made on an angled block and cut using a vibratome (Leica VT1200, Leica Microsystems, Witzlar Germany) in ice cold artificial cerebrospinal fluid (aCSF) (in mM): KCl 3, NaCl 125, MgCl_2_ 1, NaHCO_3_ 26, NaH_2_PO_4_ 1.25, glucose 10, CaCl_2_ 2 and 400 μM L-ascorbic acid, with osmolarity at 300-305 mOsm, equilibrated with 95% O2-5% CO_2_. Slices were incubated at RT until needed.

### Electrophysiology

Whole cell patch current and voltage recordings from pyramidal neurons in layer 5 somatosensory cortex were performed with a Multiclamp 700B amplifier (Axon Instruments, Union City, CA). Signals were filtered at 2 kHz and sampled at 10 kHz with Digidata 1322A (Molecular Devices, Sunnyvale, CA). Patch pipettes made from thin-walled borosilicate glass (outer diameter 1.5 mm, internal diameter 1.1 mm; Sutter instrument BF150-110-7.5) were pulled by a model P-97 puller (Sutter Instrument, Novato CA). For whole cell recording, the internal solution consisted of (in mM): K^+^ gluconate 130, HEPES 10, EGTA 0.2, KCL 10, MgCl_2_ 0.9, Mg_2_ATP 4, Na_2_GTP 0.3, phosphocreatine 20, pH adjusted to 7.2 with KOH). The osmolarity of the intracellular solution was 290-294 mOsm. Pipettes had resistances of 4-6 MΩ. Series resistance values were <15 MΩ. Junction potentials were not corrected. Both AMPAR- and NMDAR-mediated miniature excitatory post synaptic currents (mEPSCs) were recorded at −70 mV. We used the selective blockers, 1 μM tetrodotoxin (TTX), 10 µM Bicuculline methiodide (BMI), 50 µM D-(-)-2-Amino-5-phosphonopentanoic acid (D-AP5), and 1 μM TTX, 10 μM BMI, 10 μM 6-cyano-7-nitroquinoxaline-2 (CNQX), 0 mM Mg^2+^, 40 µM glycine to isolate AMPAR- and NMDAR-mediated currents respectively. Evoked excitatory postsynaptic currents were obtained by electro-stimulation of 0.1-0.2 ms duration of varying amplitude and were applied once per second with a bipolar extracellular electrode (FHC) placed at the layer 6 white matter boundary to stimulate ascending cortical inputs. Evoked AMPAR and NMDAR responses were recorded at −70 mV and +40 mV, respectively. A pCLAMP 9.2 (Axon Instruments) was used to generate stimuli and for data display, acquisition and storage, and offline analysis of mEPSCs was performed using MiniAnalysis (synaptosoft).

### UP state recording and analysis

Male mice were used at 5-6 weeks old (as described above in slice preparation). Animals were decapitated under isoflurane anesthesia and the brain quickly removed and submerged in ice cold oxygenated cutting buffer containing (in mM): NaCl 87, KCl 3, NaH_2_PO_4_ 1.25, NaHCO_3_ 26, MgCl_2_ 7, CaCl_2_ 0.5, D-glucose 20, sucrose 75, ascorbic acid 1.3, with 95% O2–5% CO2. Thalamocortical slices (400 μm) were made on an angled block (57) using a vibratome, and transferred immediately to a prechamber (BSC-PC, Warner Instruments) and allowed to recover for 1h in ACSF at 32 °C containing the following (in mM): NaCl 126, KCl 3, NaH_2_PO_4_ 1.25, NaHCO_3_ 26, MgCl_2_ 2, CaCl_2_ 2, and D-glucose 25. Then a single slice was transferred to the recording chamber and perfused with a modified ACSF that mimics physiological ionic concentrations *in vivo* which contained (in mM): NaCl 126, KCl 5, NaH_2_PO_4_ 1.25, NaHCO_3_ 26, MgCl_2_ 1, CaCl_2_ 1, and D-glucose 25, for 45-60 min before recording. For UP state analysis, spontaneously generated UP states were recorded using a borosilicate glass recording electrode (∼1 MΩ) filled with the modified ACSF and positioned in layer 4 of the somatosensory cortex.

The upward shape of UP states in our recording in comparison to the biphasic shape of UP states in previous studies (32, 37, 38) that used the same modified ACSF is due to the difference in the resistance (∼1 MΩ vs. ∼0.5 MΩ) of the recording electrode used, as shown in Fig. S2. A pCLAMP 9.2 software was used for data display, acquisition and analysis. 10 min of spontaneous activity was collected from each slice. Recordings were amplified 500×, sampled at 2.5 KHz and filtered at 2 KHz. The threshold for detection was set at 2-3X the root mean square noise. An event was defined as an UP state when its amplitude remained above the threshold for at least 200 ms (32). The end of the UP state was determined when the amplitude decreased below threshold for >600 ms (32). Two events occurring within 600 ms of one another were grouped as a single UP state (32).

### Preparation of cell suspension and fluorescent-activated cell sorting (FACS)

Mice (P40) were deeply anesthetized with ketamine (100 mg/kg) + xylazine (10 mg/kg) in saline by i.p. injection and perfused intracardially with Hanks Buffered Salt Solution (HBSS) (Thermo Scientific). The brain cortices were immediately dissected in cold HBSS buffer and cut into small pieces. Cell suspension was prepared by following the manufacturer’s instructions in the neural tissue dissociation kit (Miltenyi Biotech, Auburn, CA). Briefly, small pieces of tissue were treated with papain enzymatic mix (37 °C, 15 min) and then digested with DNase I (37 °C, 10 min), followed by careful trituration. Cell mixtures were then filtered through a cell strainer (70 μm) and resuspended in cold HBSS (5–10 x 10^6^ cells/mL) for FACS. Cells were sorted by using MoFlo MLS high-speed cell sorter (Beckman Coulter) with Summit version 4.3 software at the Tufts FACS facility.

### Immunoblot

FAC sorted astroglia were homogenized and the total protein was determined by Bradford protein assay. A total of 20 μg of cell lysate was loaded on 4 –15% gradient SDS-PAGE gels. Separated proteins were transferred onto a PVDF membrane (0.22μm, Bio-Rad) at constant 2.5Amp for 10 min using Bio-Rad Trans-Blot Turbo Transfer System. The membrane was blocked with 3% nonfat milk in TBST (Tris buffer saline with 0.1% Tween 20) then incubated with the anti-FMRP antibody (1:20; monoclonal antibody 2F5, Developmental Studies Hybridoma Bank) overnight at 4°C. On the following day, the membrane was incubated with HRP-conjugated goat anti-rabbit secondary antibody (1:5000) diluted in TBST for 1 hour at RT. Bands were visualized by ECL Plus chemiluminescent substrate using a Bio-Rad ChemiDoc MP imaging system.

### Open field test

The open field test was performed with mice (P27-P28 age) between 10:00AM and 3:00PM. We used the SmartFrame Open Field System, a Plexiglas box (41 cm x 41 cm x 38 cm) enclosing a square arena, in which mice were allowed to explore for 10 minutes. Movements were automatically detected by a grid of infrared photobeams inside the box. Parameters such as the distance traveled, and time spent in the center of the field were recorded by the Hamilton-Kinder MotorMonitor software. The amount of locomotor activity was calculated as the sum of the distances traveled in both the center and periphery regions of the arena. After each session, the box was cleaned with 70% ethanol to remove odor cues.

### Three-chamber sociability and social novelty (memory) test

Mice (P42-P48 age) were tested for social interaction behavior in a standard three-chamber plexiglass apparatus. The right- and left-side chambers contained small wire cages. This test consists of three 10-minute trials: habituation, social interest, and social novelty. For each trial, the test mouse was placed in the center chamber with free access to both the right- and left-side chambers. Data was obtained using the EthovVision video tracking software, which automatically records the amount of time the mouse spent in the right- or left-side chamber during each trial. Data from the habituation trial, where only the test mouse was in the apparatus and the wire cages were empty, served to reveal potential biases in preference for either side chamber. For the social interest trial, a male wild-type “stranger mouse” (C57Bl/6, age P21-P30) was randomly placed in either the right or left chamber under the wire cage for the duration of the trial, with the other chamber’s wire cage left empty. For the social novelty trial, a second stranger male mouse (novel stranger) of the same strain/age as the first stranger mouse (familiar stranger) was placed in the previously empty side chamber under the wire cage, with the familiar stranger remaining in its chamber from the social interest trial. An intact social memory was defined as significantly more time spent with the novel stranger compared to the familiar stranger. All mice were returned to their home cages at the end of the test. Between trials, the apparatus was cleaned with 70% alcohol to eliminate odor cues.

### Inhibitory avoidance (IA) test

Inhibitory avoidance testing was conducted using the GEMINI Avoidance System (San Diego Instruments, San Diego, CA). The apparatus consisted of two enclosed chambers with metal grid floors, separated by an automated door. In the first session (0 hr), the mouse (age P44-P59 on Day 1 of testing) was placed in one chamber of the apparatus for a 90 s habituation period, during which the chamber remained dark. At the end of the habituation period, the compartment was brightly lit and the gate to the adjacent dark compartment automatically opened. Latency was defined as the time it took for the mouse to fully enter (all 4 paws) into the dark chamber. Upon entry to the dark compartment, the door automatically closed to block the entryway between the compartments, and a single electric shock (0.5 mA, 2 s) was delivered through the floor grid to the mouse in the dark chamber. The mice were removed from the dark chamber 15 s after receiving the foot shock and were then placed back in their home cage. Subsequent trials at 6, 24, and 48 hours after the initial trial consisted of a test phase and an extinction training phase. In these subsequent trials, mice were placed in the same starting chamber for a 90 s habituation period in the dark. The chamber was then brightly lit and the gate opened, and latency to cross to the dark chamber was again measured (test phase). Once the mice crossed completely (4 paws) into the dark chamber, the gate was closed, and the mice were left in the dark chamber and allowed to explore for 200 s without an electric shock (extinction training phase). The apparatus was cleaned between trials with 70% alcohol to eliminate odor cues.

### Statistical analysis

All the resulting raw data were graphed, and statistical analyses were performed in GraphPad Prism 8 software (La Jolla, CA, USA). Data are presented as mean ± SEM. The corresponding statistical analysis is described in the legend of each figure. Significant differences were determined using single or multiple two-tailed unpaired t-test or one-way ANOVA test with post-hoc Tukey’s test, or Mann-Whitney test.

## Acknowledgements

We thank Dr. David Nelson (Baylor College of Medicine) for providing Fmr1^f/y^ and Fmr1^loxP-neo/y^ mice. Leona Tu for organizing the open field and social interaction data and Vanessa Promes for genotyping astroglial Fmr1 cKO and cON mice. We thank Dr. Andrew Tarr for help with the behavior testing. All behavior tests were performed in the Tufts University Center for Neuroscience (CNR) core facility. This work was supported by NIH grant MH106490 (Y.Y.) and FRAXA postdoc fellowship (Y.M.).

## Supplementary Information for

**Figure S1.**
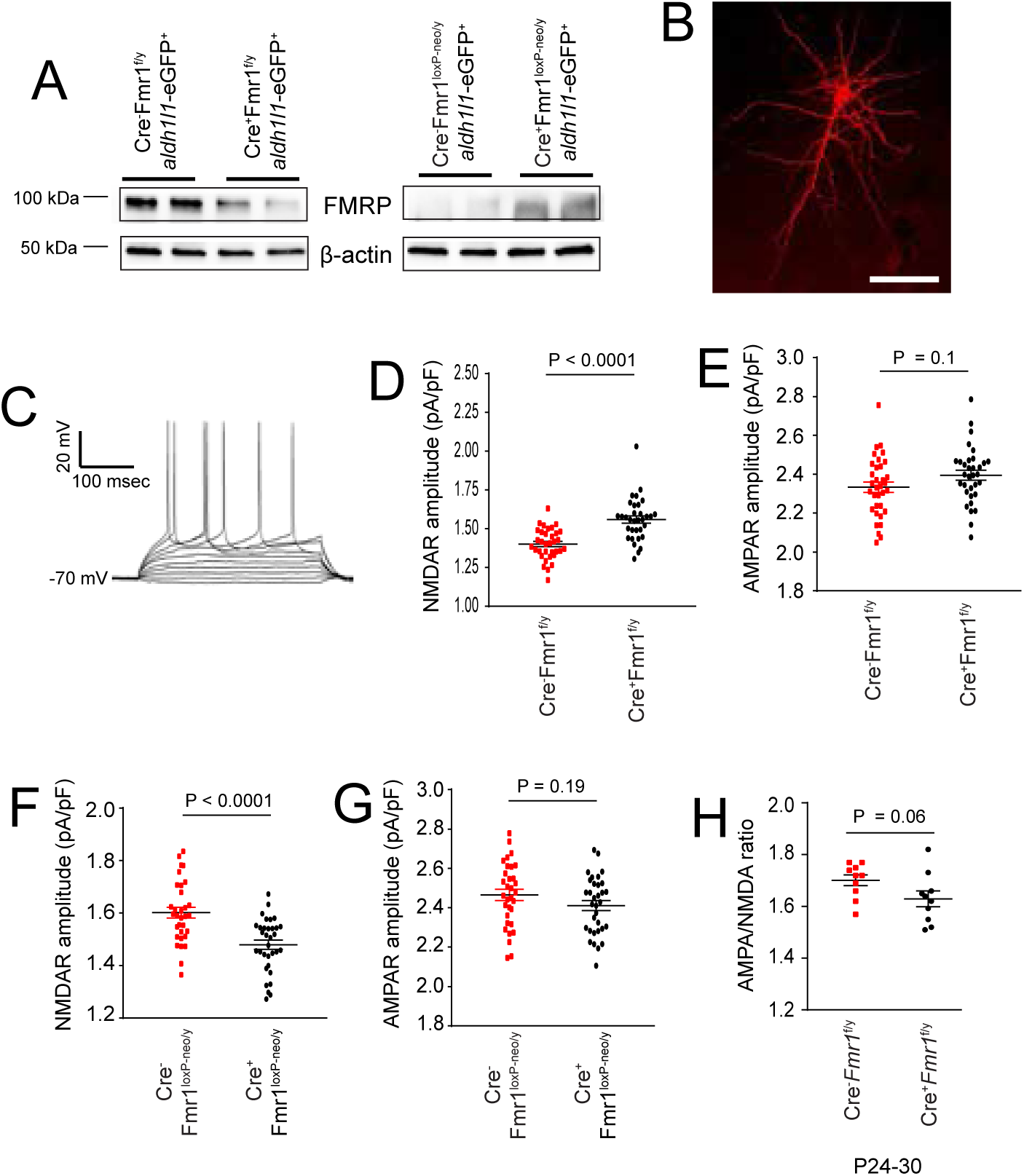
Recordings of spontaneous NMDAR and AMPAR mEPSC and evoked responses from Fmr1 KO and astroglial Fmr1 cKO and cON mice. **A.** Representative immunoblot of FMRP protein from FAC sorted cortical astroglia of astroglial Fmr1 cKO and cON mice. Representative patched pyramidal neuron (**B**) and their firing patterns (evoked by depolarizing current injection) (**C**) from the cortical layer 5 of cortical slices. Alexa-568 dye was used in filling the neuron. Scale bar: 100μm; Amplitude of NMDAR (**D**) and AMPAR (**E**) responses of Cre^-^Fmr1^f/y^ and Cre^+^Fmr1^f/y^ cortical pyramidal neurons; N=33-34 neurons from 6-8 mice per group. Amplitude of NMDAR (**F**) and AMPAR (**G**) responses of Cre^-^Fmr1^loxP-neo/y^ and Cre^+^Fmr1^loxP-neo/y^ mice; N=32-33 neurons from 6-8 mice per group. **H**. The average AMPA/NMDA ratio of cortical neurons from younger age (P24-30) Cre^-^Fmr1^f/y^ and Cre^+^Fmr1^f/y^ mice. N=10 neurons from 3-4 mice per group, All P values determined using the Mann-Whitney test.

**Figure S2.**
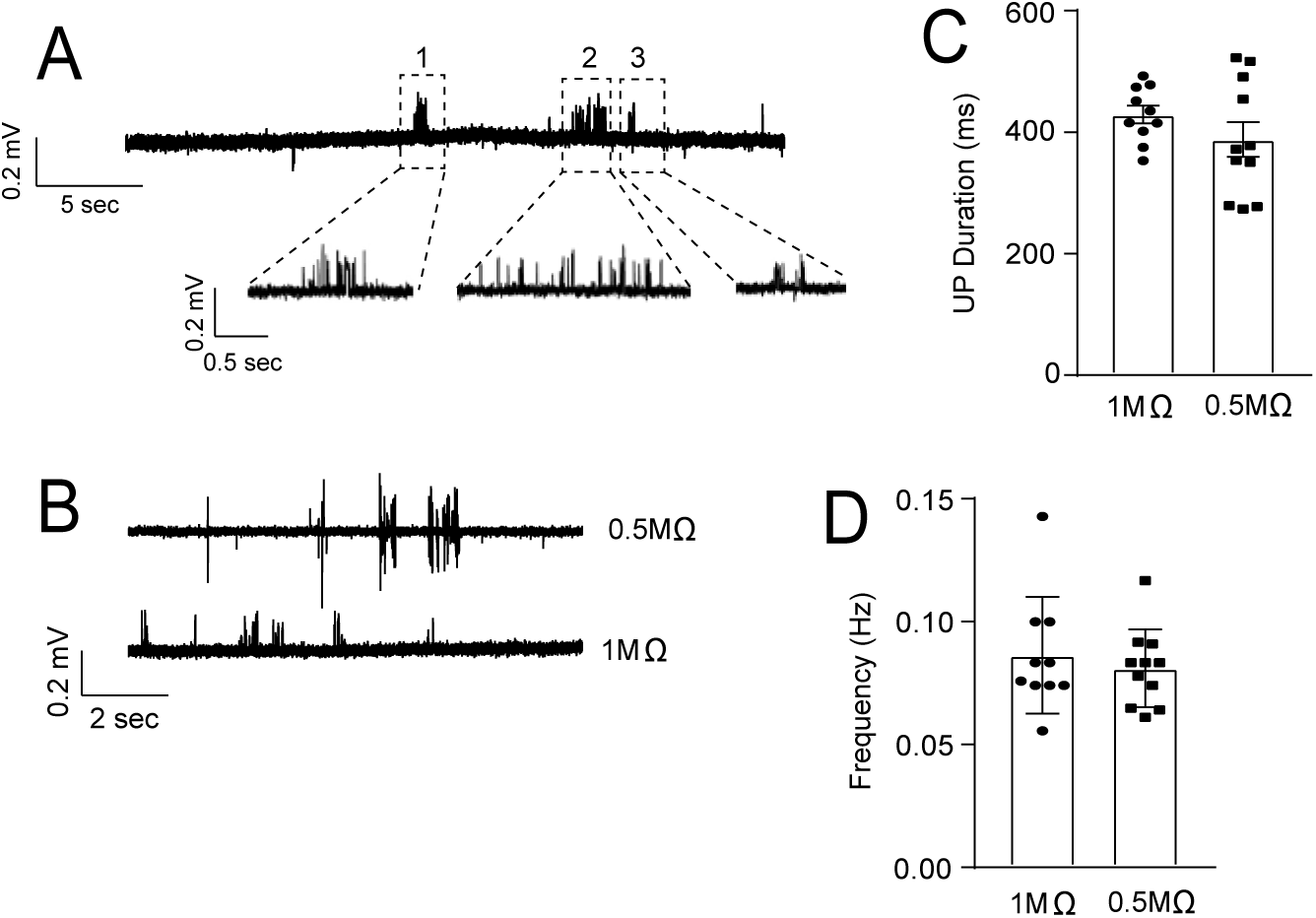
UP state recording from recording electrodes with different resistance (tip size) **A.** Representative expanded traces of UP state recording from cortical slices; **B.** Representative biphasic and upward shapes of UP states recorded using a borosilicate glass recording electrode with resistance either ∼0.5 MΩ or ∼1 MΩ respectively filled with the same modified ACSF from the somatosensory cortical slice. The duration (**C**) and frequency (**D**) of UP states recorded with a borosilicate glass recording electrode with resistance either ∼0.5 MΩ or ∼1 MΩ. N=10-11 slices per group. ms: millisecond; P values were determined using unpaired two-tailed t-test.

**Figure S3.**
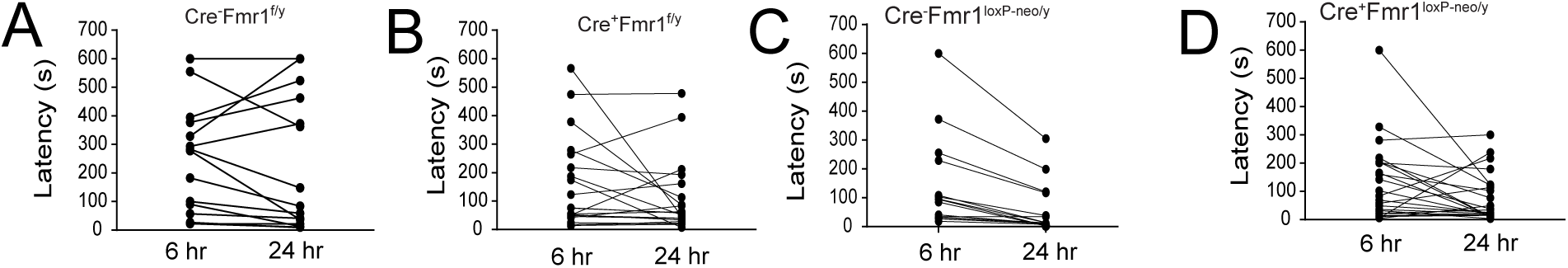
Individual mouse latency changes for the extinction phase (24h) of IA test in the astroglial Fmr1 cKO or cON mice. Latency changes from 6h to 24h of individual Cre^-^Fmr1^f/y^ (**A**) and Cre^+^Fmr1^f/y^ (**B**) mice. N=15-21 mice per group; Latency changes from 6h to 24h of individual Cre^-^Fmr1^loxP-neo/y^(**C**) and Cre^+^Fmr1^loxP-neo/y^ (**D**) mice. N=15-22 mice per group; s: seconds;

